# Previous motor actions outweigh sensory information in sensorimotor learning

**DOI:** 10.1101/2020.12.18.423405

**Authors:** Barbara Feulner, Danilo Postin, Caspar M. Schwiedrzik, Arezoo Pooresmaeili

## Abstract

Humans can use their previous experience in form of statistical priors to improve decisions. It is however unclear how such priors are learned and represented. Importantly, it has remained elusive whether prior learning is independent of the sensorimotor system involved in the learning process or not, as both modality-specific and modality-general learning have been reported in the past. Here, we used a saccadic eye movement task to probe the learning and representation of a spatial prior across a few trials. In this task, learning occurs in an unsupervised manner and through encountering trial-by-trial visual hints drawn from a distribution centered on the target location. Using a model-comparison approach, we found that participants’ prior knowledge is largely represented in the form of their previous motor actions, with minimal influence from the previously seen visual hints. By using two different motor contexts for response (looking either at the estimated target location, or exactly opposite to it), we could further compare whether prior experience obtained in one motor context can be transferred to the other. Although learning curves were highly similar, and participants seemed to use the same strategy for both response types, they could not transfer their knowledge between contexts, as performance and confidence ratings dropped to naïve levels after a switch of the required response. Together, our results suggest that humans preferably use the internal representations of their previous motor actions, rather than past incoming sensory information, to form statistical sensorimotor priors on the timescale of a few trials.

## Introduction

We often have to make decisions based on sparse and uncertain sensory information. Previous research has shown that in these cases humans use Bayesian inference where the current sensory information (likelihood) and the previously acquired knowledge (priors) are integrated, each weighted by their respective uncertainty [1,2]. While the majority of previous studies have examined whether the perceptual and sensorimotor decisions follow the rules of a Bayesian framework [1,3], less emphasis has been placed on understanding how likelihoods and especially statistical priors are learned and represented in the first place. A number of elegant recent studies have tried to bridge this gap by investigating how people learn likelihoods [4] and priors [5] to perform Bayesian computations. Interestingly, the timescale of the two types of learning varied vastly, with fast learning of likelihood but slow learning of prior distributions. It remains unknown why such an asymmetry should exist, as theoretically both types of learning are equivalent. It has been hypothesized that learning about the likelihood versus learning about the prior involves different neural mechanisms, potentially hinting to the fact that their respective distributions might be represented in different regions of the brain [6].

Learning of statistical priors is itself not a uniform process as it shows dependencies on the specific context where the learning occurs. In Bayesian framework, priors are a form of abstract knowledge [7–9], which can be generalized across different contexts. However, previous findings regarding the generalization of statistical priors have been mixed. Some studies have shown that statistical learning of priors is very narrow-band and context/modality-specific in perceptual [10] and sensorimotor domains [11,12], thus preventing learned information to transfer to different contexts/modalities. Other studies, on the other hand, provided evidence for generalization [4,13], although generalization, in some instances, seemed to occur differently for different parameters of a statistical distribution; e.g. the mean and the variance of a distribution [14]. The finding that some aspects of learning could generalize, while others could not, was confirmed by a recent study showing that, for instance, in Bayesian time estimation, priors can be generalized across stimuli, but not motor actions [15].

Therefore, despite an increasing number of studies testing generalization and transfer, the exact rules determining generalizability remain unclear. One potential reason for these seemingly contradictory results is a lack of a formal definition of *what* is learned. It has been argued that when learning is not generalized, a *policy* (i.e. a specific rule for action) rather than *knowledge* (i.e. abstract and context-independent information) is acquired through learning [16]. However, it is not clear what features of the learning dynamics determine whether a *policy* or *knowledge* is acquired during encounters with the learned information.

To investigate learning and generalizability of a prior distribution, we employed a saccadic eye movement task similar to the design of a previous study [17], where participants had to learn to locate a hidden target. The location of the hidden target corresponded to the mean of a circular normal distribution. In each trial, a visual hint sampled from the underlying distribution was shown and participants indicated their current estimate with a saccadic eye movement, looking either towards (pro-saccade) or to the exact opposite direction of the target (anti-saccade). To successfully estimate the hidden target location, participants had to combine information across multiple trials. This design allowed us to investigate whether participants formed their prior knowledge by combining the visual information or by combining the previous motor actions across time, under different saccadic response contexts. Our results from two experiments indicate that sensorimotor learning of a spatial prior in both response contexts is largely guided by previous motor plans, rather than by previous sensory input in form of visual hints. Despite the high degree of similarity of pro- and anti-saccades in their learning behavior, suggesting a motor-independent learning algorithm, the learned prior in one context did not generalize to the other. We propose that the lack of transfer between the two contexts is a natural consequence of their shared learning algorithm in which previous motor actions outweigh sensory information.

## Results

To investigate the dynamics of sensorimotor learning of a spatial prior and its dependence on the response modality, we designed an experiment where participants had to find a hidden target and indicate their guess by either a pro- or an anti-saccade. Participants learned the location of each hidden target within twenty trials, of which a block of ten trials required pro-saccades and the other block of ten trials required anti-saccades as the response modality (**Fig.1 A**). This design allowed us to probe whether the learning dynamics shows dependencies on the response modality, hence being modality-specific. We also used another task, referred to as the calibration task, to estimate each participant’s motoric noise during the visually driven execution of pro- or anti-saccades. In contrast to the calibration task, in the hidden target task the main error source is the uncertainty regarding the hidden target location. As the same motor system is used in both the hidden target task as well as the calibration task, we assumed that the motor noise affecting participants’ performance in both tasks is equal. As the motor noise in the calibration task is not time-dependent, we assumed that all time-dependent performance improvement during the hidden target task reflected statistical learning, defined as the reduction in the uncertainty regarding the location of the hidden target.

**Figure 1:**
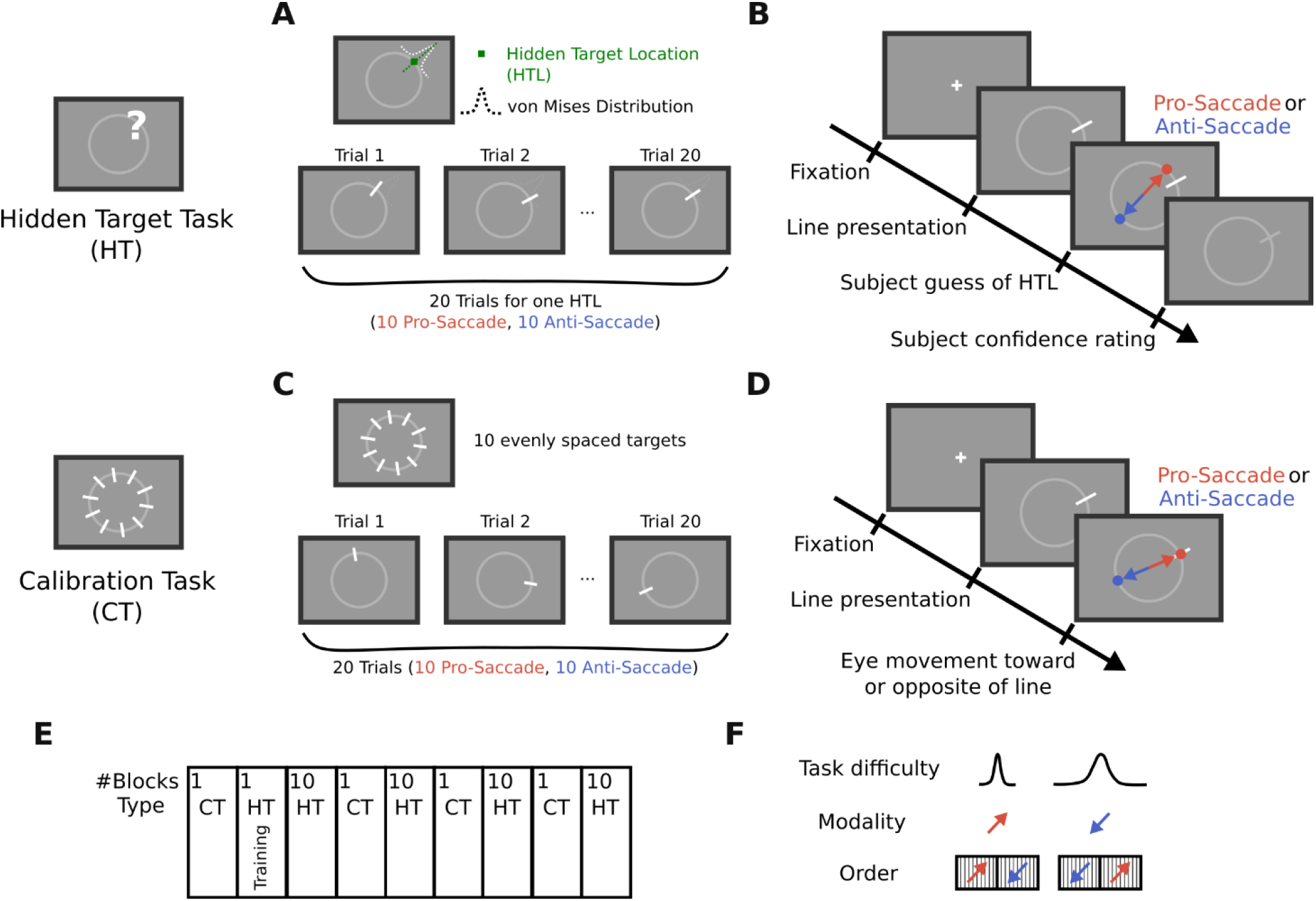
Experimental design of the hidden target task employed to study the statistical learning of a spatial prior in two different visuo-motor contexts. (A-B) Main task of the experiment. (A) Participants were told to estimate the location of a ‘hidden treasure’ on a ring by observing and combining information provided by the visual hints across trials. The hidden target location was defined as the mean of a von Mises distribution and the hints, presented at each trial, were samples drawn from this underlying distribution. Participants had twenty trials to estimate the location of the hidden target, after which a new hidden target had to be found. Participants used their gaze to indicate their responses. (B) Each trial started with a fixation period, after which the hint was presented, and participants had to indicate their guess about the location of the hidden target by either looking at it (pro-saccade response) or by looking exactly opposite to it (anti-saccade response). In half of the trials (i.e. consecutive 10 trials), participants had to use pro-saccades, and in the other half they used anti-saccades, with a randomized order across blocks. (C-D) Calibration task used to estimate the motoric error of each participant for pro- and anti-saccades. Participants had to directly look either at the lines (pro-saccade response) or exactly opposite to the lines (anti-saccade response). (E) Block-design of the experiment. (F) We compared learning across two levels of difficulty and two different response types. Task difficulty was varied by changing the standard deviation of the von Mises distribution. Finally, we tested whether knowledge could be transferred from one visuo-motor context to the other. For this we also varied the order of pro-saccade and anti-saccade responses across blocks.

### Participants successfully accumulate information and learn on a short time scale

To establish that participants were in general able to learn on a short timescale, we initially focused on the first ten trials of each hidden target block (**Fig.2 B**). In this case, in all ten trials participants responded by using the same modality, either exclusively by pro-saccades, or exclusively by anti-saccades. To quantify performance, we calculated the absolute angular error between the saccade endpoint and the hidden target location (**Fig.2 A**). By comparing the average absolute angular error in the first five trials with the average absolute angular error in the last five trials, we found that most of the participants were able to improve their estimates of the target location (i.e. their guesses) during this short timescale (paired t-test: t=7.25, p<0.0001, N=20) (**Fig.2 C**). In line with performance, participants’ confidence about the accuracy of their guesses increased during the ten trials (paired t-test: t=-4.39, p=0.0003, N=20) (**Fig.2 D**). Additionally, the absolute angular error of participants’ guesses was lower than the absolute angular error of the visual hints, meaning that participants’ guesses were closer to the center of the von Mises distribution compared to the presented visual hints (paired t-test: t=4.92, p=0.0001, N=20) (**Fig.2 E**). This shows that participants were able to combine information across trials and thereby improve their estimates of the target location, rather than just following the current visual hint. Hence, we can conclude that participants showed some form of statistical learning during the first ten trials of the hidden target task.

**Figure 2:**
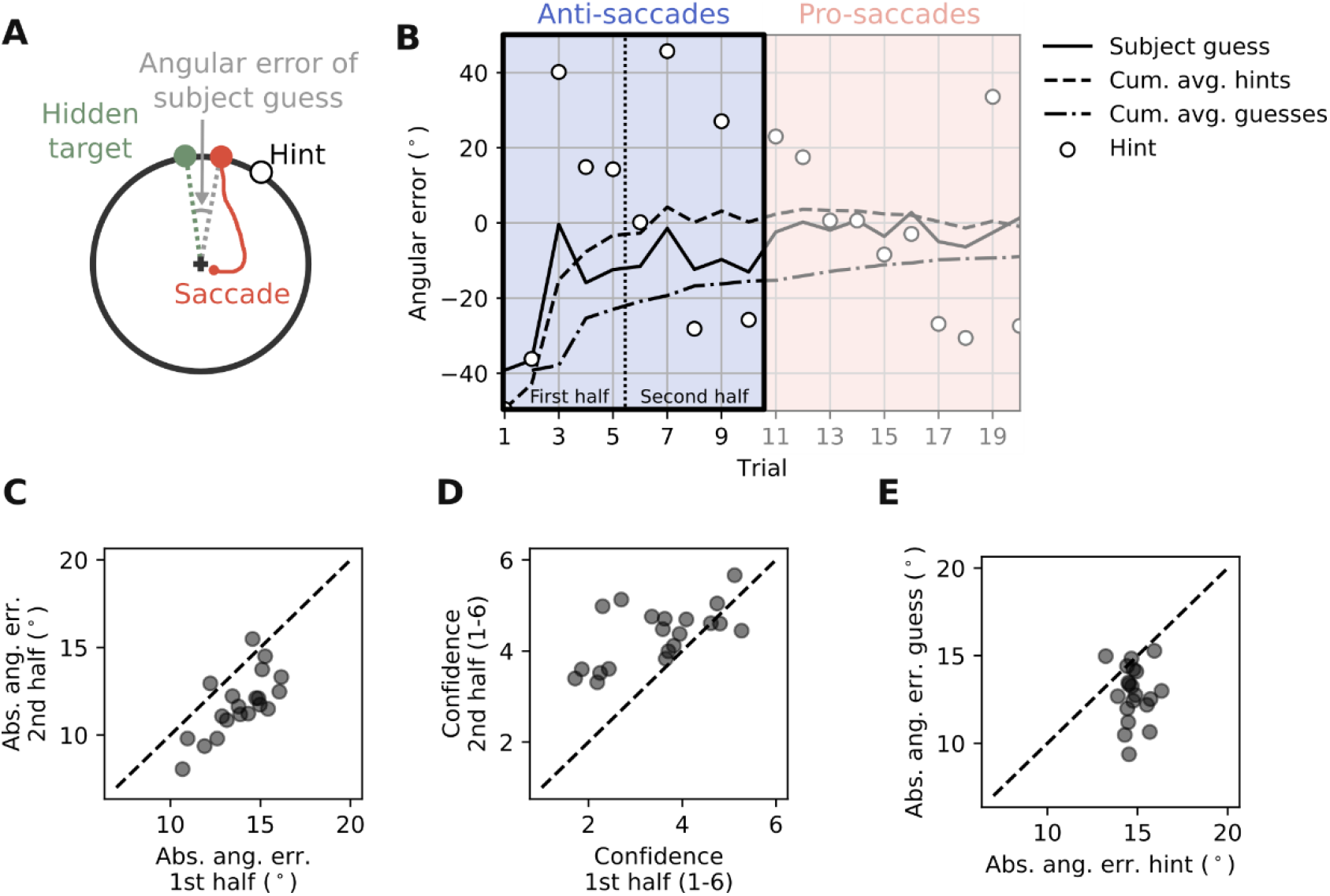
Participants successfully learn the most probable location of the target on a short time scale. (A) The angular difference between the participants’ guess and the true location of the hidden target was used to measure learning. (B) Example block. To test learning, we compared the performance in trial 1-5 (first half) to the performance in trial 6-10 (second half). (C) The absolute angular error in the second half is lower than in the first half (paired t-test: t=7.25, p<0.0001, N=20). (D) Participants’ confidence is higher in the second half than in the first half (paired t-test: t=-4.39, p=0.0003, N=20). (E) The absolute angular error of participants is lower than the absolute angular error of the visual hints, i.e., participants’ guesses are closer to the center of the von Mises distribution compared to the presented visual hints (paired t-test: t=4.92, p=0.0001, N=20).

### Learning curves are stereotypic across response modalities

To work out whether statistical learning is modality-dependent or not, we contrasted the learning performance in pro-versus anti-saccade trials in the hidden target task, as well as in the calibration task. First, we quantified the mean and the standard deviation of the respective angular error distributions, pooled across participants (**Fig.3 A&B and Table 1**). We did not find a significant bias towards a specific direction, either clockwise or counter-clockwise (one-sample t-test; mean different from zero; N=20; **Table 1**), for any of the distributions.

**Figure 3:**
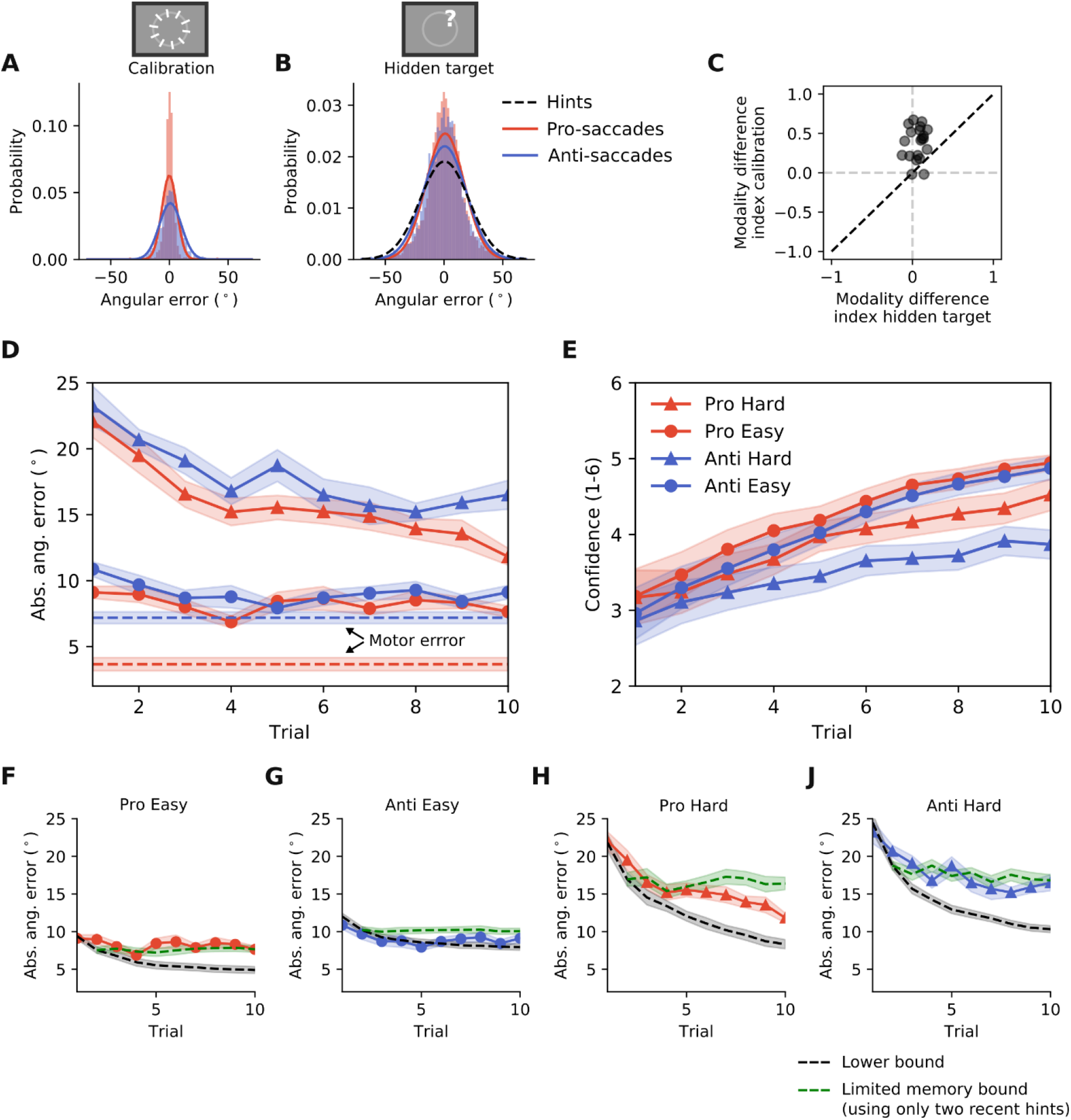
Learning curves are stereotypic across response modalities. (A) The distribution of the angular error for pro- and anti-saccade response in the calibration task. (B) The distribution of the angular error for pro- and anti-saccade response in the hidden target task. (C) Comparison of pro-/anti-saccade performance difference between the hidden target task and the calibration task. (D) Time course of the absolute angular error for each of the four different conditions (two response types x two difficulties). Here and in the following panels, except stated otherwise, shaded areas represent the standard error of the mean (N=20) (E) Time course of the confidence ratings for each of the four different conditions. (F-J) Participants’ learning curves compared to the lower bound and the limited memory bound. The lower bound is given by taking the cumulative average of all hints presented so far and adding the error due to the noise in the motor plans, estimated from the calibration task. The limited memory is similarly calculated, except that only the latest two hints are used.

**Table 1:**
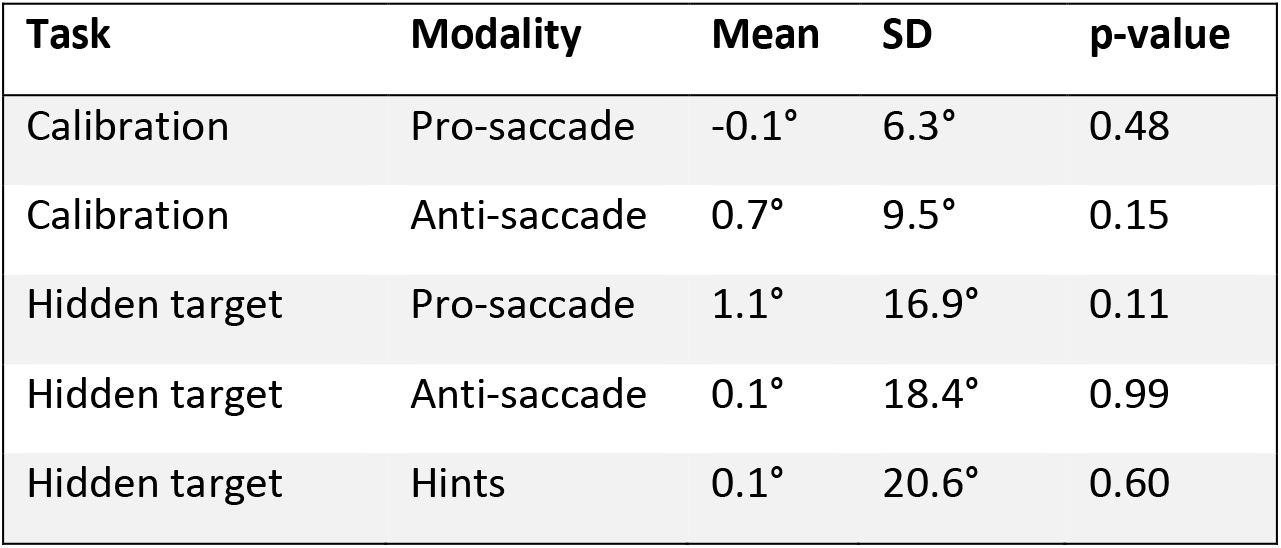
Angular error distribution for calibration and the hidden target task. Mean and standard deviation for distributions shown in Fig. 3A&B. We tested whether the mean of either of the five distributions was significantly different from zero (one-sample t-test; N=20). The corresponding p-values are shown in the last column.

| Task | Modality | Mean | SD | p-value |
| --- | --- | --- | --- | --- |
| Calibration | Pro-saccade | -0.1° | 6.3° | 0.48 |
| Calibration | Anti-saccade | 0.7° | 9.5° | 0.15 |
| Hidden target | Pro-saccade | 1.1° | 16.9° | 0.11 |
| Hidden target | Anti-saccade | 0.1° | 18.4° | 0.99 |
| Hidden target | Hints | 0.1° | 20.6° | 0.60 |

For both modalities, the standard deviation of the angular error was higher in the hidden target task than in the calibration task (pro-saccades: paired t-test, t=-11.8, p<0.0001, N=20 / anti-saccades: paired t-test, t=-10.9, p<0.0001, N=20), which is plausible as in the former task participants were more uncertain of the target location. Furthermore, the standard deviations of pro- and anti-saccades were only significantly different during the calibration task but not during the hidden target task (calibration: paired t-test, t=-3.8, p=0.0011, N=20 / hidden target task: paired t-test, t=-1.7, p=0.1108, N=20). To quantify this difference further, we calculated a modality difference index (described in Materials and Methods section), which was significantly higher in the calibration task compared to the hidden target task (paired t-test, t=-6.44, p<0.0001, N=20, **Fig.3 C**). These results provide the first evidence that statistical learning is modality-independent.

To see whether there is a difference between pro- and anti-saccades depending on the difficulty of the task, we next looked at the influence of both experimental factors (pro-/ anti-saccade response, broad/narrow hint distribution) on participants’ performance. To this end, we conducted a linear mixed model analysis of the single trial data (i.e., the absolute angular error), with *task difficulty* and *response type* as fixed effects and participant identity as random effect. Using this statistically more powerful approach, we found a main effect of task difficulty (t=20.6, p<0.0001; contrast: easy-hard=-8.28°±0.27°(SE), t=-31.2, p<0.0001), as well as response type (t=2.0, p=0.0469; contrast: pro-anti=-1.32°±0.27°(SE), t=-5.0, p<0.0001). Additionally, an interaction effect was found between the task difficulty and response type (t=2.2, p=0.0299), indicating that there is only a difference between pro- and anti-saccade responses in the hard task condition (hard_pro-hard_anti=-1.90°±0.38°(SE), t=-5.035, p<0.0001; easy_pro-easy_anti=-0.74°±0.37°(SE), t=-1.988, p=0.1927). The same pattern was found when we examined participants’ confidence ratings: participants were more confident in pro-saccade trials and in the easier task condition (main effect *response*: t=-4.2, p<0.0001, contrast: pro-anti=0.28±0.03, t=10.3, p<0.0001 / main effect *difficulty*: t=-9.25, p<0.0001, contrast: easy-hard=0.47±0.03, t=17.4, p<0.0001 / interaction effect: t=-4.4, p<0.0001). These results show that there is a small influence of the used visuo-motor modality on participants’ performance and confidence in the hidden target task, if the hint distribution is broad (hard task condition). As shown in **Fig.3 A-C**, this difference is much smaller than the difference we observed in the calibration task.

To investigate whether the response type had an influence on the statistical learning itself, we included the trial number as another fixed effect in our linear mixed model (as a continuous variable). If the response type influenced participants’ performance in a trial-independent manner, it would act as a general offset. In contrast, if statistical learning were modality-dependent, we would expect an interaction between the trial number and the response type (pro- versus anti-saccades). Visual inspection of the learning curves for pro- and anti-saccade responses suggested the former, and the main effect influencing the shape of the learning curves seemed to be the task difficulty (**Fig.3 D-E**). For participants’ performance, measured as the absolute angular error, we found a significant main effect of difficulty (t=21.4, p<0.0001) and a significant interaction between trial number and difficulty (t=-7.807, p<0.0001), but no significant interaction between trial number and response type (t=0.650, p=0.516). Further analysis of the interaction effect showed that only for blocks with high task difficulty there is a significant effect of trial number (hard: slope=-0.84±0.08, t=-10.57, p<0.0001 / easy: slope=-0.12±0.08, t=-1.50, p=0.13). This result was further supported by the analysis of the confidence ratings, as participants’ confidence time course (**Fig. S1**) was also mainly dominated by the influence of task difficulty (interaction effect *trial* x *difficulty*: t=-8.858, p<0.0001) and not by the response type (interaction effect *trial* x *response*: t=-1.006, p=0.315). Together, these results show that both performance and confidence increase in a modality-independent manner during learning of a spatial prior.

### Performance is suboptimal

Since at each point in time, participants have only seen a limited number of samples from the underlying distribution of the target location (in form of visual hints), it is theoretically impossible to correctly estimate the distribution’s mean, i.e., the hidden target location in our task. In principle, only for an indefinite number of samples, the sample mean equals the population mean. For the hidden target task, this means that participants could theoretically only reach zero absolute angular error if an infinite number of trials were seen. Taking this into account, we can compare participants’ performance to the time course of the theoretical statistical uncertainty due to seeing a limited number of samples (i.e., sampling error, see also Materials and Methods section). This provides a lower bound on participants’ uncertainty, indicated by the variance of angular errors *σ*^2^:

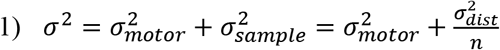

where *n* is the number of visual hints seen so far, 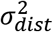 is the variance of the von Mises distribution and 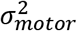 is the motoric noise measured in the calibration task. We can then calculate the theoretical optimal performance in terms of the mean absolute angular error from this optimal variance estimate (see Eq 2 in Materials and Methods section). Comparing participants’ performance to this lower bound shows that for the first trial in each hidden target block, participants perform as well as they possibly can (**Fig.3 F-J**), independent of the task difficulty or the used response modality – pro- or anti-saccades. However, as soon as they need to combine multiple samples to estimate the hidden target location, they perform suboptimal (**Fig.3 F-J**, black dashed line is optimal). We hypothesized that the reason for this suboptimal behavior is limited memory, as to perform optimally would mean to remember every single hint which has been presented so far. So instead of calculating the cumulative mean of all visual hints presented so far (which results in optimal performance), we calculated the performance estimate for an alternative strategy where subjects only average the latest two visual hints (**Fig.3 F-J** green dashed line). This limited memory estimate could describe participants’ performance qualitatively better, as it captured the shallower learning curves for both response types. In summary, the analysis of different aspects of performance did not show large differences in statistical learning between pro- and anti-saccades.

### Strategy is modality-independent

So far, we have seen that statistical learning in the pro- and in the anti-saccade context is similar in terms of the absolute performance and the shape of the learning curve. Next, we wanted to test whether participants use different strategies in the pro-versus in the anti-saccade blocks. The first candidate strategy would be to only look at the visual hints in each trial, which would mean that no learning is happening, and that participants’ behavior is only visually-driven. Instead, the optimal strategy, which would result in the lower bound we calculated before, would be to calculate the cumulative average of every hint seen so far in each trial. Besides taking into account the hints, visually presented to the participants, we can also imagine that the behavior is driven by an internal state that promotes looking close to where one has been looking before, i.e., to follow previous guesses (**Fig.4 A**). To test these different hypotheses, we fitted several single predictor linear regression models, for each proposed strategy. Through a model comparison (see Materials and Methods for the details) we found that the best single predictor model is the cumulative average of previous guesses (**Fig.4 B**). Splitting the data into pro- and anti-saccade blocks and repeating the analysis showed that the best single predictor model does not depend on the used response type (**Fig.4 C-D**). Thus, this provides another piece of evidence that the statistical learning happening in our task is independent of the response type.

**Figure 4:**
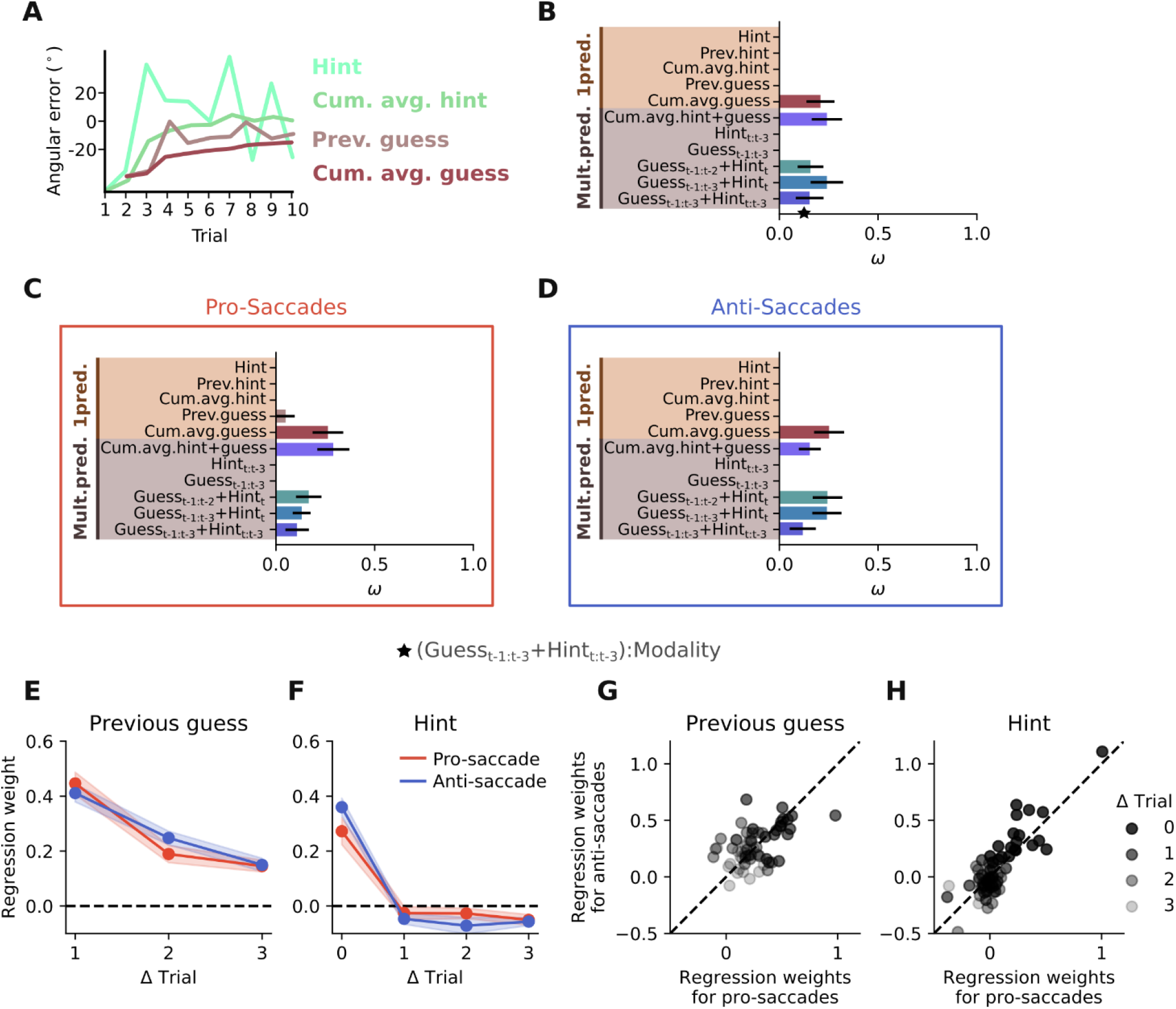
Strategy is modality-independent. (A) Different predictors used to explain the participants’ single trial estimates. (B) Model comparison between various single and multiple predictor models. Shown are the weights ω, which represent the probability that a model is the best among the ones considered. (C-D) Same as (B) but performed on two different data sets, one consisting only of pro-saccade response trials, the other consisting only of anti-saccade response trials. (E-H) Regression weights for a model including participants’ last three guesses and the current and last three visual hints. (E) Regression weights put on the last three guesses. (F) Regression weights put on the current, as well as the last three hints. (G-H) Participants put similar weight on guesses and hints in pro- and anti-saccade response trials (paired t-test; n.s.).

In a second step, we also tested several multi-predictor models to get a more detailed view on participants’ strategy. We tested multiple combinations of past guesses, and past and present visual hints. Models based on the previous three guesses or the current and previous three visual hints did not perform as well as the best single predictor model. In contrast, models which relied on combinations of external and internal information, i.e., the visual hints and previous guesses, performed as well as the best single predictor model, even if they only looked three time steps in the past (**Fig.4 B**). Again, splitting the data into pro- and anti-saccade blocks did not affect the main trend (**Fig.4 C-D**).

Lastly, to find the exact weighting participants put on their previous guesses and the current and previous hints, depending on the used response type, we fitted a model with all three previous guesses, the current and three previous hints, and the response type as predictors. We found that participants used external information only from the current trial, ignoring the hints from previous trials (**Fig.4 F**). Instead, to combine information across trials they relied on their internal estimations from the past (**Fig.4 E**). We obtained similar results when models that only included either the past guesses or the past visual hints were tested **(Fig. S5)**. Including the response type in the model allowed to estimate separate regression weights for pro- and anti-saccades. Again, we didn’t find any significant difference in the weighting participants put on their own guesses versus given hints, depending on whether they use pro- or anti-saccades for response (**Fig.4 E-H**; paired t-test: n.s.). In summary, we found that participants used the same strategy, regardless of the response type, to solve the task, which provides further evidence for the modality-independent learning hypothesis.

### Drop in performance after response switch

Despite the similarity in performance and observed strategy, hinting at a general algorithm used for statistical learning in our task, it is still possible that learning happens for each visuo-motor modality in a very specific manner and that both just look similar in terms of performance and strategy. In this case, it would not be possible to generalize between modalities. To test this, we included trial eleven until twenty where participants had to continue looking for the same hidden target (and were also instructed that all twenty trials belong to one hidden target), but had to use the other visuo-motor modality than in trial 1 until 10 (**Fig.5 A**). We considered two alternative hypotheses. The first states that participants learn in a modality-independent fashion and store the acquired knowledge in an abstract form, which allows a complete transfer of previous experience to a new response modality after a switch (**Fig.5 B**, black solid line). The second hypothesis would be that participants perform trial eleven until twenty as if there was no previous experience, which would suggest a modality-specific implementation of the learned knowledge (**Fig.5 B**, gray dashed line). To test which of these hypotheses is true, we compared the performance in trial 10 (last trial before response switch) to the performance in trial 11 (first trial after response switch). We found that the performance is significantly worse in trial 11 than trial 10 (paired t-test, t=-6.5, p<0.0001, **Fig.5 C**). The performance decrease in trial 11 was consistent across all 4 conditions (**Fig.5 F**). This shows that there is no direct transfer of knowledge across response types. After ruling out the modality-general implementation, we wanted to test if there is any measurable interaction between the first ten trials and the second ten trials. For this, we compared the starting points of each learning curve – trial 1 and trial 11 (**Fig.5 D**), as well as the end points of each learning curve – trial 10 and trial 20 (**Fig.5 E**). There was no significant difference between these, suggesting that the learning curves are identical. These results were corroborated by analyzing confidence ratings, as we also observed a drop in confidence at trial 11 compared to trial 10, in line with our results on performance (paired t-test trial 10 - trial 11, t=3.7, p=0.0016, **Fig. S1&S2**)

**Figure 5:**
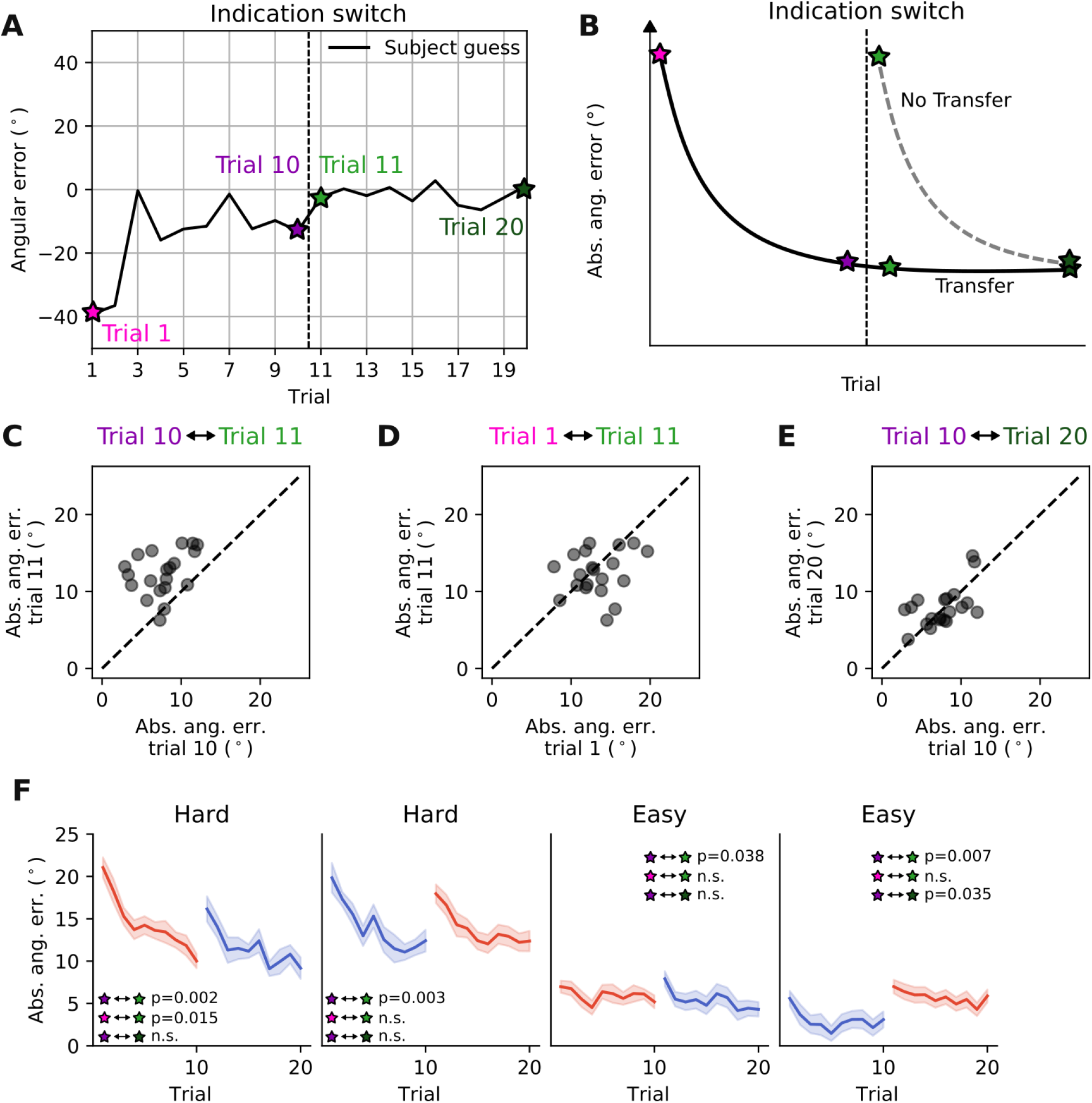
Drop in performance after response switch. (A) To test the knowledge transfer hypothesis, we analyzed all trials within a block, encompassing trials before and after the switch. We specifically focused on the difference between trial 10 (before response switch) and trial 11 (after response switch). (B) If knowledge is transferred, we expect similar performance in trial 11 as in trial 10. In contrast, if no knowledge is transferred, we expect trial 11 to show similar performance to trial 1. (C) Performance in trial 11 is worse than in trial 10 (paired t-test, t=-6.5, p<0.0001). (D) Performance in trial 11 is similar to performance in trial 1 (paired t-test; n.s.; N=20). (E) Performance in trial 10 is similar to performance in trial 20 (paired t-test; n.s.; N=20). (F) Performance time course for different difficulty levels and pro-/anti-saccade orders. To illustrate the difference due to statistical learning only, we subtracted the motor error estimated from the calibration task. Insets show the result for the same statistical test as in (C-E), but performed separately on the data of each condition.

### No knowledge transfer between visuo-motor modalities

As modeling showed that participants rely on their previous guesses rather than previously seen visual hints (**Fig.4 E**), we wished to test whether this is also the case across the time that a response switch occurs. For this, we calculated the regression weights on previous guesses in a time-dependent manner (**Fig.6 A**). If knowledge is not transferred between different visuo-motor modalities we would predict: 1) Participants highly rely on the visual hint in trial 11 and do not use previous guesses to inform their decision. 2) They are also not able to use previous guesses further in the past, if these guesses were obtained before the response switch. Our analysis showed that indeed these were the case. We found that participants’ estimates in trial 11 (after response switch) are independent of their guess in trial 10 (before response switch). In contrast, at all other time points they used previous guesses to inform their current decision (**Fig.6 B**). As participants did not use their previous experience in trial 11, we expected that they instead rely highly on the hint presented in trial 11. Regression analysis confirmed this hypothesis (**Fig.6 C**), although we found that in three out of the four conditions the weight on the current hint in trial 11 is lower than in trial 1. Only for the easy task difficulty level and the ordering of first anti-saccades and then pro-saccades, the weight on the presented hint at trial 11 was as high as at the beginning of the block (**Fig.6 C** red dashed line). Also inspired by the initial modeling results on participants’ strategies (**Fig.4**), we tested the influence of previous guesses further in the past (two trials (D) and three trials (E)). Again, we found that there is no knowledge transfer across the response switch. In summary, these results show that participants were unable to use their past estimates, made with another response type, to inform their current guess.

**Figure 6:**
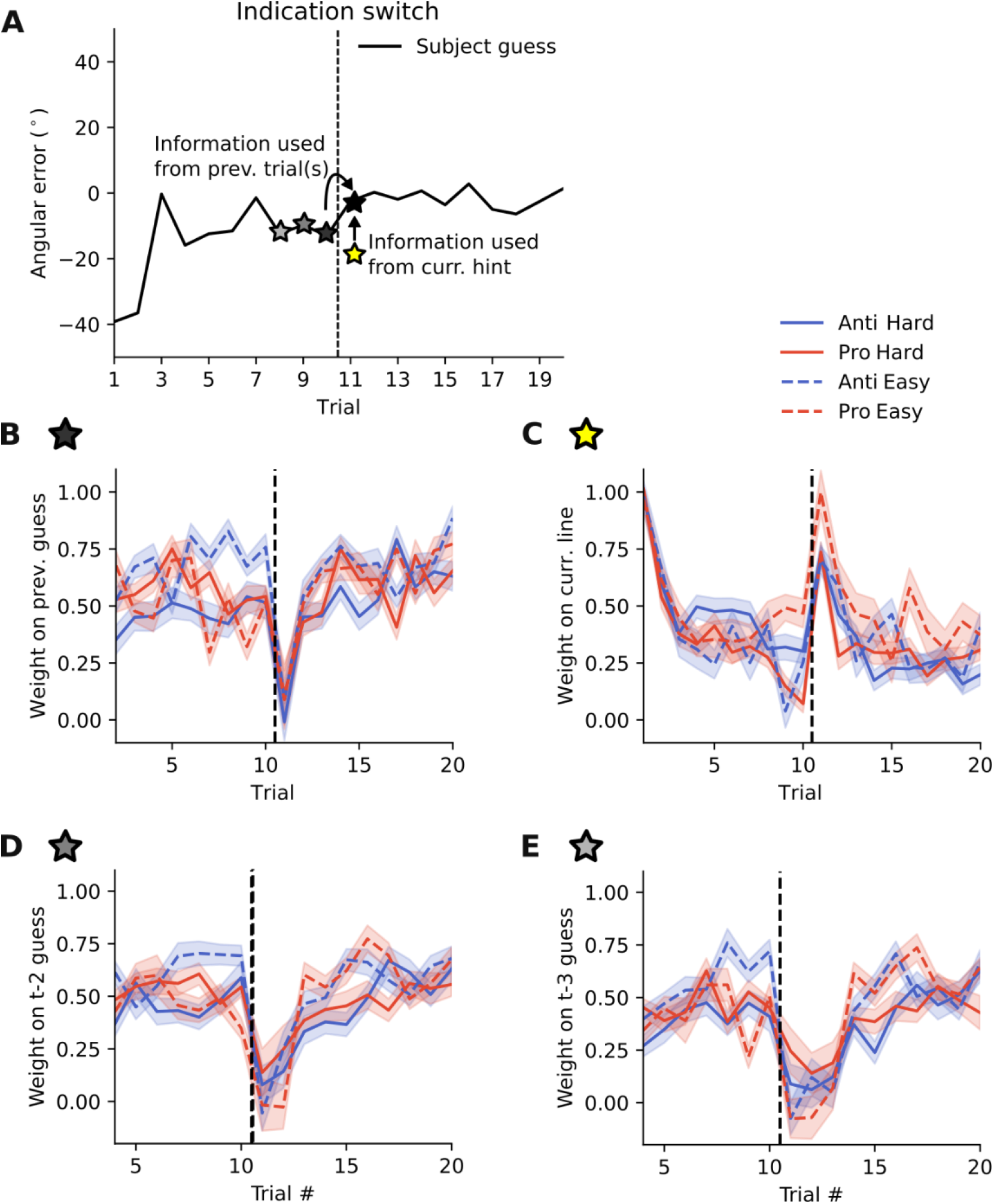
No knowledge transfer between visuo-motor modalities. (A) To test if there is any knowledge transferred from the experience with one response type to the other, we regressed the current guess against previous guesses/current hint. (B) Participants’ estimates at trial 11 (after response switch) are independent of the estimates at trials 10 (before response switch). In contrast, at every other time point, participants use previous experience to inform their current guess. (C) At trial 11, participants highly rely on the information coming from the current hint. (D-E) Besides transfer from one trial to the next (B), there is also no transfer from trials further in the past across the response switch.

One possible explanation for these results is that participants had difficulties in understanding that the hidden target location had remained the same across first and second 10 trials (or between trial 10 to 11). To rule out this possibility, we performed a second experiment with an independent set of participants (N=20) where we further emphasized that the target location was the same across the two halves of the experiment. Furthermore, we asked the participants at the end of each block whether they were aware that all twenty trials performed so far belonged to the same hidden target. We obtained overall similar results in this experiment (**Fig. S3**), as learning performance, as well as information transfer and confidence were disturbed after the response switch between trial 10 and 11, despite the fact that participants were explicitly instructed that all twenty trials belonged to the same hidden target and that they were also aware of this rule (**Fig. S3 B**). This indicates that the inability to transfer knowledge between response types is not because of a cognitive misunderstanding of the task.

## Discussion

The aim of the current study was to characterize the dynamics of prior learning and its dependence on the type of motor response used to report choices. We found that participants could learn a sensorimotor prior within a few trials, with the learning time course being independent of the response type (pro- or anti-saccades). By using a model-comparison approach, we further demonstrated that participants relied more on their own guesses from previous trials compared to visual hints provided in previous trials – again – independent of the response type. This suggests that prior knowledge is represented in terms of previous motor actions and not incoming, external information provided by the visual hints. To verify this hypothesis, we tested whether participants could generalize their learned prior knowledge from one motor context to the other – a switch from pro- to anti-saccades or vice versa. We found that switching the response type caused participants to reset to naïve levels of performance, indicating that experience from one response type could not be generalized to the other. This was the case even despite explicit instructions and participants’ awareness that pro- and anti-saccade trials belonged to the same hidden target location. Our results suggest that humans learn sensorimotor priors through monitoring their previous motor decisions rather than external sensory hints. The dependence of learning on past motor decisions discourages generalization of the learned knowledge to conditions where a different visuo-motor mapping is needed.

Our findings suggest that prior knowledge is represented in a motor specific manner during early learning, which is in line with previous studies reporting motor specific priors in different paradigms [15,16]. We could furthermore identify one potential reason for why generalization is not possible in such contexts, as we found that participants do not memorize the external information from previous trials (visual hints in our case), but instead they memorize their own actions in each trial (**Fig.4 E-F**). As our task required an estimation in every trial, indicated by either a pro- or an anti-saccade, the memory of each trial’s decision was probably represented as the motor action taken to indicate the guess. This could also explain why tasks which do not require an explicit response via a motor action are more generalizable [15]. In these cases, the memory from previous trials is potentially formed in a more abstract way, as participants only have to ‘think’ about their decision, but not perform any specific action. The finding that prior knowledge is built on internal decisions, compared to external cues, could therefore unify some previous controversial findings about prior generalization.

Despite the suggested motor specific formation of prior knowledge, the algorithm to combine previous experiences to inform the current decision seemed to be similar for both tested response contexts (**Fig.4 E-H**). This suggests that there is a general procedure for how humans combine previous experiences. However, whether prior knowledge can generalize or not depends on the specific manner through which previous experience or decisions are stored in memory (e.g., in terms of motor actions or more abstract decisions). In other words, although at an algorithmic level learning is independent of the response modality, the learned information is stored with a format that is specific for each modality. This explanation is in line with the previously proposed dissociation between learning a *policy* versus *knowledge* [16], and further narrows the space of testable predictions regarding the neural implementation of these different types of learning, as the well-described neuronal machinery of pro- and anti-saccades [18,19] could allow a characterization of how the two types of learning occur in the brain, for instance through using neuroimaging techniques.

Our study is different from previous studies as it does not test prior learning in a condition where there is also sensory uncertainty [5,17]. In these general task designs, participants are asked to perform a task trial-by-trial and are not explicitly told to combine information from previous trials. Prior learning in these cases is therefore implicit and potentially unconscious. Furthermore, learning is mostly observed by analyzing how participants combine the noisy sensory information in a given trial with the formed prior. It is therefore not directly possible to resolve which of the two is learned, as observed changes in this combination could potentially come from a changed likelihood distribution, from a changed prior distribution, or from changes in both distributions. Because of these limitations, we designed our task such that the sensory information in each trial was given by a clearly visible hint. We then explicitly asked the participants to combine the information across trials to find the hidden target location. Compared to previous prior learning studies [5], we could therefore directly look at the ability to learn statistical regularities over trials. This more general statistical learning context has also been studied previously, for example showing that learning can happen rapidly within a dozen trials if feedback is provided [20]. However, to our knowledge, pure statistical learning, without trial-by-trial feedback, together with generalization has not been studied in this context.

The two different task contexts we investigated in this work are distinct to previous studies, as we did not test generalization from one effector to another, such as performing a task with the right hand and switching to the left [11,21], or switching from a motor to a perceptual task [16]. Instead, our two contexts represent two distinct cue-action mappings, though performed with the same modality (oculomotor system). By cue-action mapping we mean that participants had to indicate their guess (the internal cue) with two different response types – pro- and anti-saccades (the actions), dependent on the task context. Potentially, generalization could be easier between modalities compared to generalization between different cue-action mappings. Given the specific design of our task, we cannot differentiate whether participants form a motor independent spatial prior, which is aligned to the given cue-action mapping, or whether they form their prior directly at the motor level. In both cases generalization would fail, matching our experimental observation. Potentially, participants learn a spatial representation in the pro-saccade context, where the correct estimate lies close to their performed motor action endpoint. Then, in the anti-saccade context, they don’t follow this estimate and solely invert their motor plan, but instead they form a ‘pro’ representation of the hidden target location in this new context, where again the performed motor action endpoint is close to the acquired spatial estimate. In this interpretation, anti-saccades are not really anti-saccades, but pro-saccades relative to the participants’ estimates and only visual information is inverted. What speaks for this interpretation is the fact that participants seemed to be closer to the optimal learning performance in the anti-saccade condition (**Fig.3 F-G**), although this was only the case when the visual hints were narrowly distribution.

Our setup allowed us to simultaneously evaluate participants’ performance, as well as their confidence in their given estimation. Interestingly, confidence also decreased after the response type switch, suggesting that participants were aware that they cannot generalize between both contexts. On the other hand, a control experiment, where we explicitly asked participants to indicate after each block whether they were aware that both response type contexts belonged to the same hidden target estimation, showed that that they knew that information could be combined across both contexts (**Fig. S3 B**). Together, these results suggest that participants’ inability to transfer knowledge from one response context to the other was not due to conscious misunderstanding, but more likely due to the specific mechanisms of how the prior is formed unconsciously.

Although our results, especially the inability to generalize across different motor contexts, suggest that statistical learning is implemented in a low-level, motor-dependent way (**Fig.5&6**), there might be potentially alternative explanations for why we could not see generalization in our experiment. In the following we will discuss these alternative interpretations. One potential explanation for the lack of generalization could be an interference between the internal memory of the target location, which is disturbed by the information slide, presented between trial 10 and 11 indicating response switch (**Supplementary Video 1**). Another possibility is that the short timescale of only ten trials is not enough to form a general representation of the estimated target location. Potentially, initial learning is motor-dependent, but over time this is transformed to a motor-independent knowledge, which can then be transferred to other motor contexts. Another possibility is that participants would need to train on our task for more than one session, such that they can learn to adapt their way of forming the prior to something which allows for generalization across response contexts. Finally, one major reason for a motor-context dependent learning could be that we did not provide external feedback to guide the learning process (unsupervised learning). Instead, participants had to rely solely on their internal feedback, potentially coming from the motor system. Future studies will be needed to test these possible explanations.

Spatially directed movements such as saccadic eye movements and reaching are an integral part of our daily activities and acquired skills (e.g., imagine a cellist or a tennis player). Both types of movements are profoundly influenced by statistical priors [1,22]. Furthermore, saccadic eye movements provide detailed sensory information about a scene and are tightly linked to the allocation of attention, hence being instrumental for active vision [23,24]. Typically, the effect of statistical priors on saccadic eye movements is investigated by using a fixed set of potential target locations, where the probability of target appearing in some locations (hence the uncertainty of that location) is more than the others. The typical finding of these studies [22,25] is that saccade latencies became shorter with increasing prior probability of the corresponding target location. One novel aspect of the current study is to test how spatial priors can be learned through eye movements under more complex settings where the possible target locations are not fixed, and probabilities could only be inferred through tracking noisy information over time. In comparison to the saccadic eye movement, reaching movements have enjoyed a more rigorous characterization of the learning dynamics [5,12,14,16]. Inspired by these studies, we characterized the dynamics of statistical learning through eye movements, that are a more accessible motor plan to be tested in the laboratory, and the learned information acquired through their execution can directly impact the very way that the brain samples the sensory information [26].

## Materials and Methods

In this study, we report the results of two experiments investigating how a spatial prior is learned under different oculomotor response contexts. The second experiment was identical to the first and served as a control for ensuring that participants were aware of how the location of the learned prior varied across blocks of the experiment (see the description of the Experimental design). We therefore, describe the methods common to both experiments and mention the difference where they apply.

### Participants

In total, 41 participants were recruited for this study. 21 participants (age range 22-38 years, M=26.05, SD=3.80, 10 females) took part in the first experiment. All participants had normal (N=9) or corrected-to-normal vision (N=12). One participant was excluded from the data of the first experiment as the post-experiment questionnaire indicated that the participant had misunderstood the task. 20 participants (age range 23-38 years, M=28.95, SD=5.07, 11 females) took part in the second experiment (with no exclusion) and all had normal (N=16) or corrected-to-normal vision (N=4). Participants were recruited from the general population of the city of Göttingen, Germany, using flyer and online advertisement and received cash financial compensation for their participation. Participation was voluntary; all participants were informed about the study procedure and gave written consent prior to the test session. The study was approved by the local ethics committee of the “Universitätsmedizin Göttingen” (UMG), under the proposal number 15/7/15.

### Experimental setup

The stimuli were presented at the center of a calibrated ViewPixx/EEG monitor (VPixx Technologies, QC Canada, dimension: 53 x 30 cm, refresh rate: 120 Hz) with a resolution of 1920 x 1080 pixels at a viewing distance of 60 cm. All experiments were scripted in MATLAB, using Psychophysics toolbox [27]. Eye movements were measured using the Eyelink1000+ eye tracking system (SR Research, Ontario, Canada) in a desktop mount configuration, recording the right eye, with a sampling rate of 1000 Hz. A chin rest was used to stabilize the participant’s head. The EyeLink camera was controlled by the EyeLink toolbox in MATLAB [28]. At the beginning of each experiment, as well as after every 10 blocks of the *Hidden Target Task* (see the Experimental design), the eye tracking system was calibrated using a 13-point standard EyeLink calibration procedure. Calibration was repeated until an average error of maximum 0.5 visual degrees was achieved and the error of all points was below 1. If the calibration accuracy dropped during the experiment, e.g., due to the subjects’ movement, the experimenter recalibrated the eye tracking system again.

#### Experimental design

The experiment comprised two tasks: a calibration task (to estimate the motor error of pro- and anti-saccades) and the main task, referred to as the ‘hidden target task’ (**Fig.1**). There were in total four blocks of the calibration task (n=20 trials in each block) and forty blocks of the hidden target task (n=20 trials in each block). Each experiment started with a block of the calibration task followed by one training block for the hidden target task (n=10 trials in this block). The data from this training phase was not analyzed. Thereafter, the experiment proceeded to the main task where participants performed 10 blocks of the hidden target task followed by 1 block of the calibration task. This sequence was repeated 4 times (**Fig.1 E**).

In a second experiment, we used the exact same experimental design but enforced the instruction that the location of the hidden target remained the same across a block of 20 trials although the response type changed halfway through (from anti- to pro-saccade or vice versa). For this, we adapted the information slides shown during the experiment (**Supplementary Video 1&2**). Furthermore, we asked the participants after every twenty trials of a hidden target whether they were aware that the last twenty trials belonged to the same hidden target. Participants then had to press a button to indicate their response, either yes or no.

#### Hidden Target Task

Participants were instructed to look for a ‘hidden treasure’ location on a ring with a radius of 7.5°, centered in the middle of the screen (**Fig.1 A**). The word ‘hidden treasure’ was used in our instructions to the participants to make the task more realistic and engaging, however we will refer to the task as the ‘hidden target’ task throughout. Each trial started with a fixation period, where participants had to fixate for 0.5 s on the white cross (size = 0.1875°, color: white, displayed on a half-grey background) in the middle of the screen (**Fig.1 B**). After that, a white line (length =1.125, color: white) was presented and participants had 3 s to estimate the hidden target location for this trial and indicate their guess either by looking at it (pro-saccade) or by looking opposite of it (anti-saccade) and fixate their estimated location for 0.5 s. Thereafter, participants rated their level of confidence in their guess on an discrete scale from 1 to 6, where 1 means very uncertain and 6 means very certain about the target location. The confidence rating had to be done within 4 seconds.

Participants were told that they had twenty trials to guess the location of the hidden target, after which a new hidden target had to be found. To estimate the hidden target location participants had to closely monitor the location of a line that was presented in every trial and served as a visual hint. The hidden target location was the mean of a von Mises distribution and each visual hint was a sample drawn from this distribution (see below) [17]. Hence, by paying attention to the location of the hint across trials, participants were able to infer the underlying distribution of the hidden target location. Participants indicated their estimates by looking either at where they thought the hidden target was located on the ring (pro-saccade), or at a location directly opposite to it (anti-saccade). Ten consecutive trials of a block of twenty trials required pro-saccade responses, and the other ten anti-saccade responses. The type of the required response (either pro- or anti-saccade) was visually indicated by an instruction display presented every ten trials. Participants were instructed to perform the same type of response for ten trials in a row, until the response type changed.

In total, there were forty hidden target blocks. The target location of each block, which is the mean of the von Mises distribution, was randomly drawn from a fixed set of twenty locations evenly distributed on the circle. Thus, each location only appeared twice during the experiment. To familiarize the participants with the connection between the hints and the hidden target location, ten training trials were performed in the beginning of each experiment. After the ten training trials, participants saw all the ten lines together on the screen, as well as the correct hidden target location. Furthermore, after each hidden target block, participants saw where the actual hidden target location was, but they did not receive feedback about their performance on a trial-by trial basis. As such, in our experiments learning was unsupervised.

Four different experimental conditions, counter-balanced across blocks, were tested. Each block was either *easy* or *hard*, controlled by adapting the concentration of the von Mises distribution, and either ordered with first pro-saccade then anti-saccade, or first anti-saccade then pro-saccade response (**Fig.1 F**). For the easy task condition the concentration of the von Mises distribution (defined by κ which is a measure of dispersion, where 1/κ is equivalent to the variance σ^2^_dist_ of the distribution) was 30 (σ_dist_~10°), for the hard task condition it was 5 (σ_dist_~26°) and for the training it was 80 (σ_dist_~6°). As the concentrations of these distributions are relatively large, we could treat the von Mises distribution as a normal distribution and use standard statistics.

#### Calibration Task

The aim of this task was to quantify the participant-specific motor error of the visually-driven pro- and anti-saccades. Each block of this task consisted of twenty trials, from which the first ten were pro-saccades and the last ten were anti-saccades. On each trial, one out of ten equally distributed locations on the circle (same circle as in the hidden target task) were selected and a target line was presented at that location (**Fig.1 C**). Participants were instructed to look either directly at the displayed line (pro-saccade in the first ten trials), or directly opposite to where it appeared (anti-saccades, second ten trials). Additionally, it was highlighted that this task is completely independent of the hidden target task. Each trial consisted of an initial fixation phase, where participants had to fixate on the white cross in the middle of the screen for 0.5 s. After that, a white line appeared and participants had to either look at it or opposite of it (**Fig.1 D**). In the beginning of a block of ten trials, participants received an instruction display indicating whether they had to perform pro- or anti-saccades during the upcoming trials.

#### Successful response

For both tasks, a successful response was defined as follows. Participants had to move their gaze from the central fixation point towards a peripheral location on the ring. As soon as they moved away from the fixation and crossed a circular threshold of 5.375° a successful response was possible. To complete the response, participants furthermore had to fixate on one specific point on the screen, by holding their gaze for 0.5 s within an area with a radius of 1.

### Analysis

#### Data pre-processing

The recorded raw eye movement data was transformed to MATLAB files by using a MATLAB library for eye movement analysis [29]. Participants’ estimates in each trial were calculated offline by averaging the eye movement data of the last 100 ms, out of the total 500 ms necessary for a successful response. The main eye movement parameter that we analyzed was the angular distance between the participant’s estimate and the true hidden target location (**Fig.2 A**). Failed trials were excluded from the analysis. A trial could fail in several ways. Firstly, there could have been a disturbance with the eye tracking system or the participant’s calibration so that the gaze position was not correctly detected, which made it necessary to re-calibrate the eye tracker. Secondly, the participant could have been too slow to indicate their guess in time (3 s). Thirdly, to exclude erroneous pro-instead of anti-saccades and vice versa, we analyzed the distribution of angular errors and found a bimodal distribution (**Fig. S4**). We set a threshold at 100°, which separated both modes (**Fig. S4**). Thus, every trial with an absolute angular error bigger than 100° was categorized as failed because of the wrong response type.

### Data analysis

Statistical analyses were done using R and Python. To evaluate learning we used paired, two-sided t-tests to compare several parameters within a participant. We did not use circular statistics as subjects’ responses where highly localized on the ring (**Fig.2 A-B & S4**). Learning was assessed by measuring the decrease in the absolute angular error between the participant’s estimate and the true hidden target location across trials. Given the assumptions that the participant’s estimates are normally distributed and centered around zero, the mean absolute angular error can be related to the variance of the Gaussian distribution by:

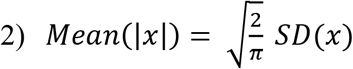

To compare the performance difference between pro- and anti-saccades, we calculated a ‘modality difference index’ (**Fig.3 C**). For this, we first calculated the median of the absolute angular error for pro-saccade and anti-saccade response trials. The modality difference index is then given by the difference between the two medians, divided by their sum. We started our analysis by only using the data from the first ten trials of each block (**Fig.1–4**). Only when we looked at the transfer between modalities, we used all twenty trials of each block (**Fig.5–7**).

#### Theoretical bounds for learning performance

To evaluate participants’ performance, we computed the theoretical lower bounds on the absolute angular error they could potentially achieve by using all the information that was available to them. Participants could use the previously seen visual hints and their memory of their previous motor actions (referred to as the previous guess) to infer the most probable location of the target on each trial. These sources of information are error-prone since on each trial participants had only seen a limited number of visual hints (i.e., sampling error) and their previous responses contained motoric noise. We assumed that these two sources of error are independent. The error due to the limited number of visual hints was calculated as the cumulative mean of all hints seen so far, which represents an optimal way of combining samples over time to estimate the mean of a distribution. A constant motor error, measured for each participant during the calibration task, was used to represent the motoric noise. The variance of the joint estimate, derived from combining visual hints and motor actions, was then calculated by adding the variance of the two sources, based on the assumption of their independence (cf. Eq.1). In addition, we also calculated a suboptimal, ‘limited memory’ lower bound to account for the information loss across time. This model makes use of the latest two visual hints, instead of all, to infer the statistical distribution of the target. To understand how participants used these sources of information to make decisions on a trial-by-trial basis, we employed a detailed model-comparison approach as described below.

#### Modeling participants’ behavior

We used linear regression models to analyze the single subject behavior. Four potential strategies we wanted to test were: 1) looking directly at the visual hints in each trial, 2) estimating the cumulative average of all visual hints so far, 3) looking at the same location as in the previous trial, 4) estimating the cumulative average of all previous guesses (as given by their saccadic responses) (**Fig.4 A**). Strategy (2), calculating the cumulative average of all hints seen so far is directly linked to the lower bound on performance described above. This is an optimal statistical strategy to combine all observed hints and therefore produces the optimal performance. In principle, strategy (2) is also equivalent to a Bayesian optimal strategy, as in our case each hint has objectively equal certainty and should therefore be weighed equally.

We started with the four described single predictor models to see which of the four mentioned strategies best describes participant’s behavior. The dependent variable was the angular error of a participant’s estimate in a certain trial. The independent variable was the angular error of the estimate, given by one of the above-mentioned strategies. Only the data from trial 4 onward was included to test all models on the same data, since we wanted to test the influence of up to three trials in the past on the current trial. In a second stage, we tested linear regression models with multiple independent variables. We included previous guesses from up to three time steps in the past, as well as the visual hints from the current and up to three time steps in the past. Each of the described regression models was fitted to the single subject data. To compare these models, we calculated BIC, delta BIC, and Bayesian weights [30], to assess the likelihood of each model being the best fit to the data (cf. Evaluating model performance). Since these values are normalized, they can be used to determine the model that on average best fits the participants’ data.

#### Evaluating model performance

To compare the different linear regression models presented above we used a model comparison evaluation based on Bayesian weights [30]. For this, we firstly calculated BIC values for each model. Each BIC value was rescaled by calculating ΔBCI, which is calculating the difference to the smallest BIC value in the group of models considered. This forces the best model to have ΔBCI=0 and the other models to have positive values. We then calculated Bayesian weights ω with:

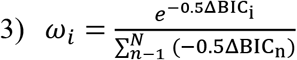

The Bayesian weights of all tested models in **Fig.4 B** sum up to 1 and define the probability of being the best model, among the one tested.

## Acknowledgements

We thank Tomke Schoss for her help with the pilot study. This work was initially supported by a seed fund grant from Leibniz Science Campus Primate Cognition, Göttingen, Germany to CMS and AP and continued by an ERC Starting Grant (no: 716846) to AP. CMS was supported by an Emmy Noether Grant from the German Research Foundation (SCHW1683/2-1).

## Authors’ contribution

BF and AP conceptualized the project. BF, CMS and AP designed the task. BF implemented the experiment. DP collected the data. BF analyzed the data. BF and AP interpreted the results and wrote the first draft of the manuscript. All authors revised the manuscript. AP and CMS acquired funding.

## Supplementary figures

**Figure S1:**
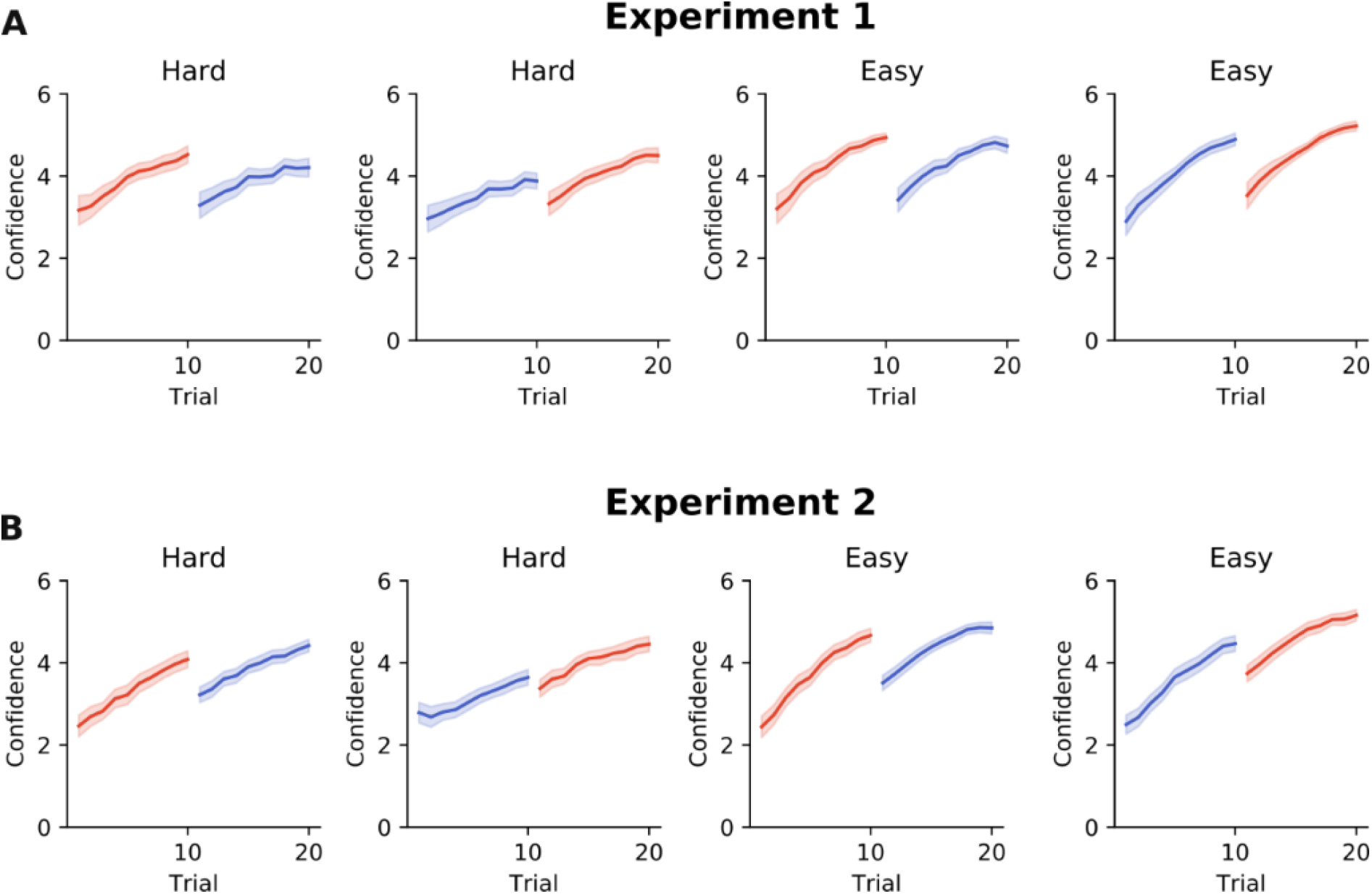
Confidence time course.

**Figure S2:**
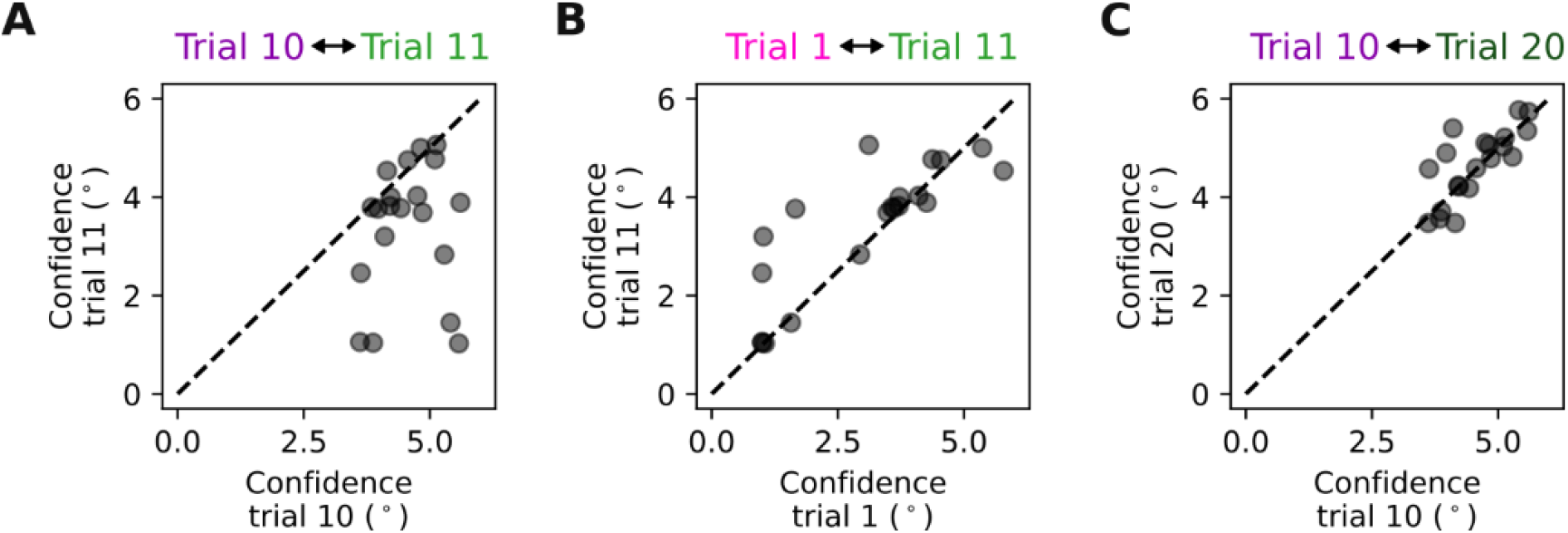
Confidence drops after response switch. Figure S1 and S2 are supplementary to Fig.5 in the main text. The confidence ratings demonstrated the same results as observed by analysing the absolute angular errors of the eye movements, as there was a decrement in confidence between trials 10 and 11 as shown in S1-A) for Experiment 1, as well as in S1-B) for Experiment 2, for all experimental conditions. The drop in confidence between trial 10 and 11 was significant (S2-A). The confidence of trial 11 was not different from trial 1 (S2-B), and the last trial of a block of 20 trials was not significantly better than trial 10 (S2-C). These results support the observation that after a switch in response modality, learning starts from a naïve level.

**Figure S3:**
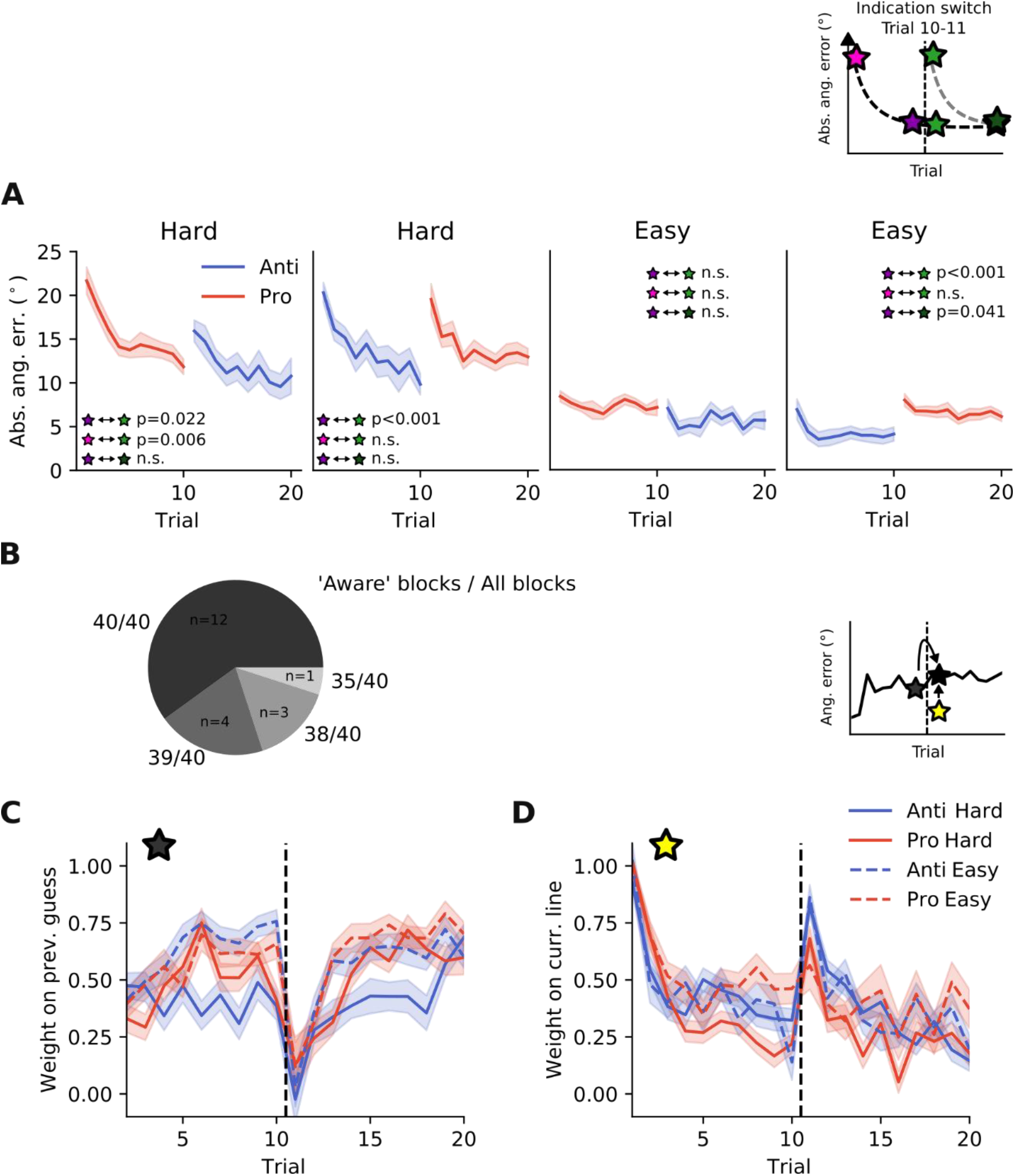
Second experiment with reinforced instructions shows similar results. This figure is supplementary to Fig.5 and Fig.6 in the main text. These results demonstrate that we obtained similar results, when participants were explicitly instructed that the location of the hidden target remains the same after a switch. Additionally, participants had to report whether they were aware of this rule, thus reinforcing the instructions. A) Also in this experiment, performance dropped to naïve levels after a switch in response type. B) The majority of participants reported to be aware of the rule. C) The weighting of previous guesses dropped between trial 10 and 11 (when the switch occurred) and D) instead more weight was put on visual hints.

**Figure S4:**
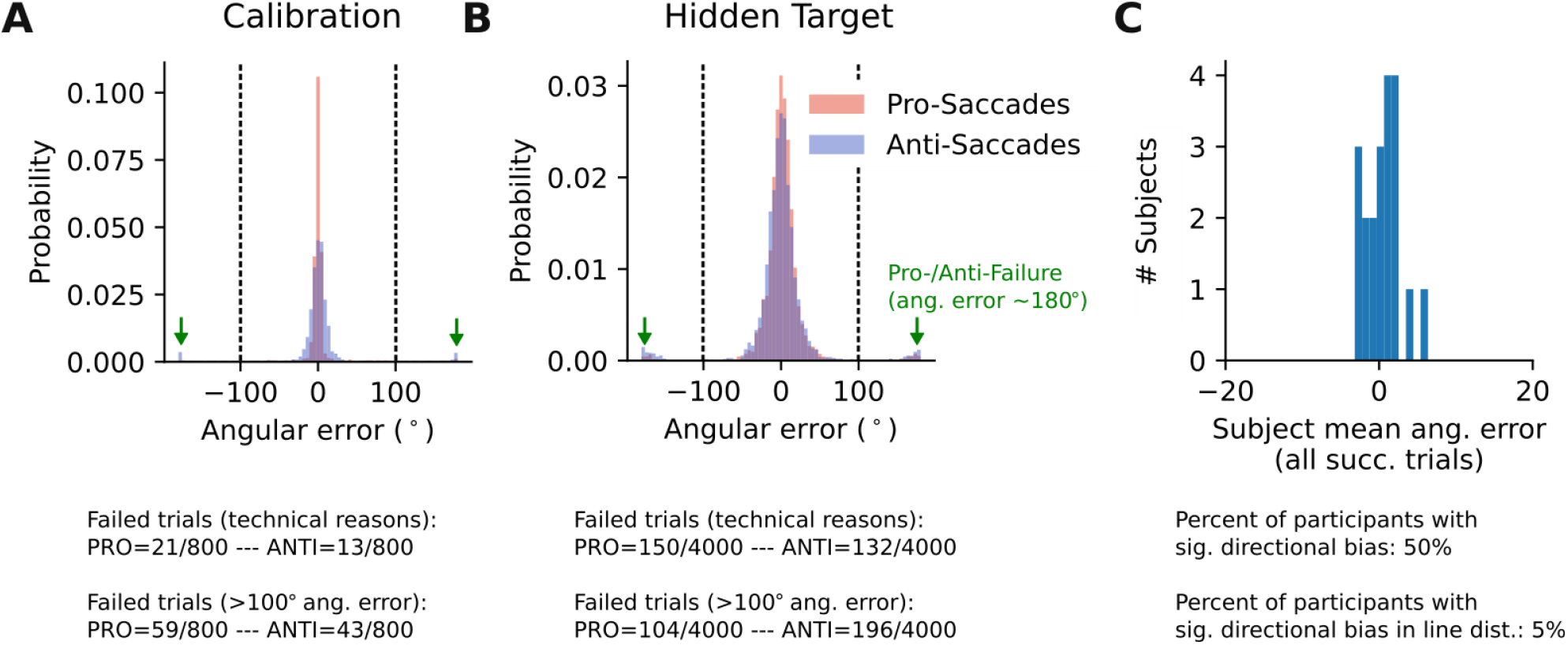
Threshold for failed pro-/anti-saccades and subject-wise directional bias. This figure is supplementary to Fig.3 in the main text. It demonstrates the rationale for excluding erroneous pro- and anti-saccades. A) Data of Calibration Task. B) Data of Hidden Target task. In both panels, subjects’ estimates are shown without excluding ‘wrong’ saccades due to pro-/anti-saccade error. As shown in A and B, there is a bimodal distribution of angular errors, especially in the hidden target task. The peaks at ±180° error represents the failed pro- /anti-saccades, meaning trials where pro-saccades should have been made but the subject responded with anti-saccades and vice versa. We set a threshold at 100 degree to distinguish correctly aimed pro- or anti-saccades. C) Distribution of the mean angular error across participants indicates that there was no directional (CCW or CW) bias.

**Figure S6:**
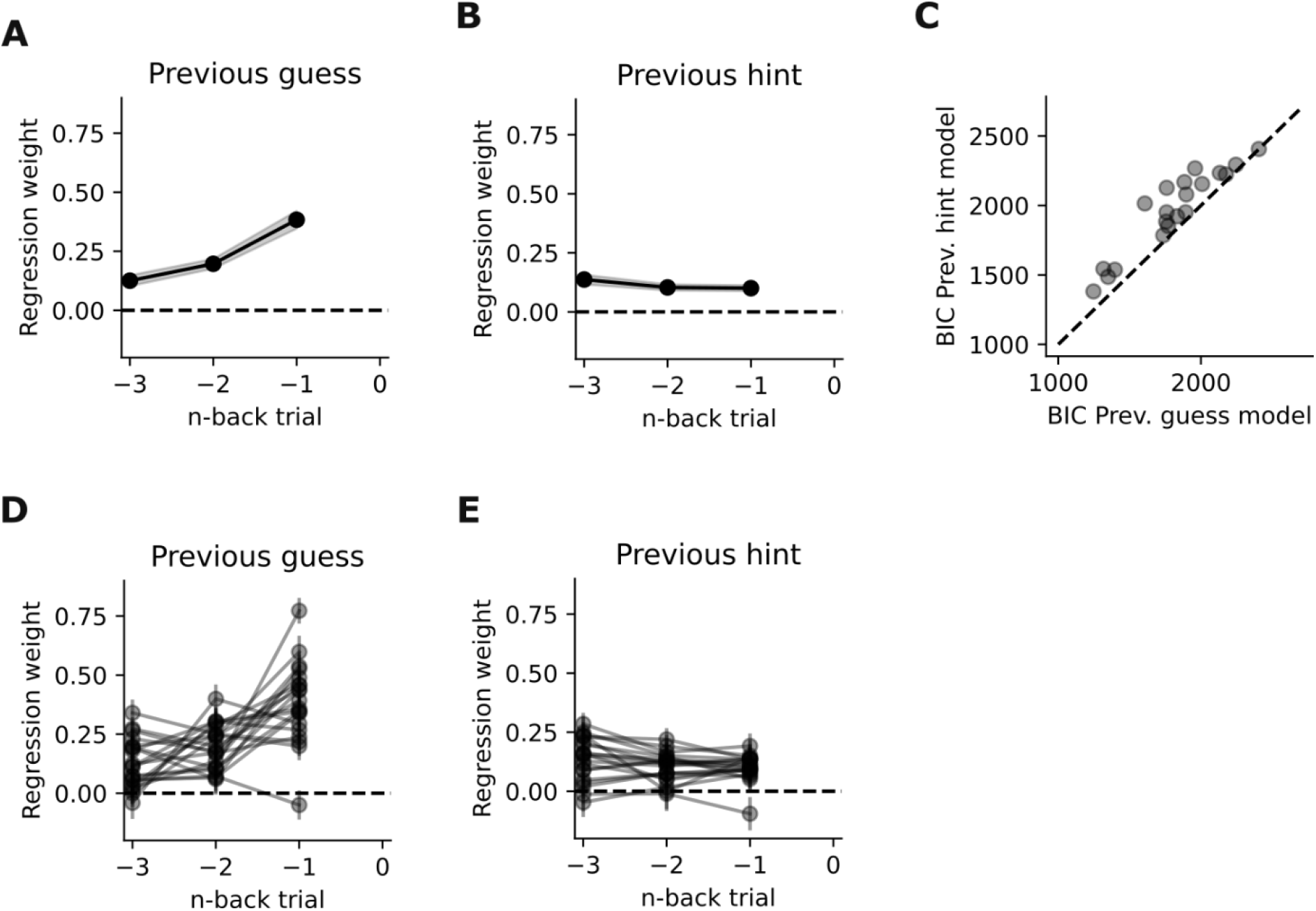
Memory traces for previous guesses and hints. This figure is supplementary to Fig.4 in the main text. Here, we quantified how much weight is put on previous hints or guesses, tested in separate models each including either only the guesses or only the visual hints. A) Regression weights of the model including previous guesses as a predictor B) Same as A for a separate model including visual hints as a predictor 3) comparison of the model shown in A against the model shown in B. It can be seen that the previous guess models consistently outperformed the visual information/hints models, confirming the finding that subjects’ behavior is better predicted by previous actions than external visual information D&E) Same as A&B, but showing the single subject regression weights instead of the average weights.

## References

1. Körding KP, Wolpert DM. Bayesian integration in sensorimotor learning. Nature. 2004;427: 244–247. doi:10.1038/nature02169

2. Wei K, Körding K. Uncertainty of feedback and state estimation determines the speed of motor adaptation. Front Comput Neurosci. 2010;4: 1–9. doi:10.3389/fncom.2010.00011

3. Ernst MO, Banks MS. Humans integrate visual and haptic information in a statistically optimal fashion. Nature. 2002;415: 429–433. doi:10.1038/415429a

4. Sato Y, Kording KP. How much to trust the senses: Likelihood learning. J Vis. 2014;14. doi:10.1167/14.13.13

5. Berniker M, Voss M, Kording K. Learning Priors for Bayesian Computations in the Nervous System. Brezina V, editor. PLoS One. 2010;5: e12686. doi:10.1371/journal.pone.0012686

6. Vilares I, Howard JD, Fernandes HL, Gottfried JA, Kording KP. Differential representations of prior and likelihood uncertainty in the human brain. Curr Biol. 2012;22: 1641–1648. doi:10.1016/j.cub.2012.07.010

7. Battaglia PW, Hamrick JB, Tenenbaum JB. Simulation as an engine of physical scene understanding. Proc Natl Acad Sci U S A. 2013;110: 18327–18332. doi:10.1073/pnas.1306572110

8. Tenenbaum JB, Kemp C, Griffiths TL, Goodman ND. How to grow a mind: Statistics, structure, and abstraction. Science. Science; 2011. pp. 1279–1285. doi:10.1126/science.1192788

9. Tenenbaum JB, Griffiths TL, Kemp C. Theory-based Bayesian models of inductive learning and reasoning. Trends Cogn Sci. 2006;10: 309–318. doi:10.1016/j.tics.2006.05.009

10. Frost R, Armstrong BC, Siegelman N, Christiansen MH. Domain generality versus modality specificity: The paradox of statistical learning. Trends in Cognitive Sciences. Elsevier Ltd; 2015. pp. 117–125. doi:10.1016/j.tics.2014.12.010

11. Hewitson CL, Sowman PF, Kaplan DM. Interlimb generalization of learned bayesian visuomotor prior occurs in extrinsic coordinates. eNeuro. 2018;5. doi:10.1523/ENEURO.0183-18.2018

12. Yin C, Wang H, Wei K, Körding KP. Sensorimotor priors are effector dependent. J Neurophysiol. 2019;122: 389–397. doi:10.1152/jn.00228.2018

13. Kiryakova RK, Aston S, Beierholm UR, Nardini M. Bayesian transfer in a complex spatial localization task. J Vis. 2020;20: 17–18. doi:10.1167/JOV.20.6.17

14. Fernandes HL, Stevenson IH, Vilares I, Kording KP. The generalization of prior uncertainty during reaching. J Neurosci. 2014;34: 11470–11484. doi:10.1523/JNEUROSCI.3882-13.2014

15. Roach NW, McGraw P V., Whitaker DJ, Heron J. Generalization of prior information for rapid Bayesian time estimation. Proc Natl Acad Sci U S A. 2017;114: 412–417. doi:10.1073/pnas.1610706114

16. Chambers C, Fernandes H, Kording KP. Policies or knowledge: Priors differ between a perceptual and sensorimotor task. J Neurophysiol. 2019;121: 2267–2275. doi:10.1152/jn.00035.2018

17. Dekleva BM, Ramkumar P, Wanda PA, Kording KP, Miller LE. Uncertainty leads to persistent effects on reach representations in dorsal premotor cortex. Elife. 2016;5. doi:10.7554/eLife.14316

18. Munoz DP, Everling S. Look away: The anti-saccade task and the voluntary control of eye movement. Nature Reviews Neuroscience. Nature Publishing Group; 2004. pp. 218–228. doi:10.1038/nrn1345

19. Ford KA, Goltz HC, Brown MRG, Everling S. Neural processes associated with anti-saccade task performance investigated with event-related fMRI. J Neurophysiol. 2005;94: 429–440. doi:10.1152/jn.00471.2004

20. Chukoskie L, Snider J, Mozer MC, Krauzlis RJ, Sejnowski TJ. Learning where to look for a hidden target. Proc Natl Acad Sci U S A. 2013;110: 10438–10445. doi:10.1073/pnas.1301216110

21. Kumar A, Panthi G, Divakar R, Mutha PK. Mechanistic determinants of effector-independent motor memory encoding. Proc Natl Acad Sci U S A. 2020;117: 17338–17347. doi:10.1073/pnas.2001179117

22. Brodersen KH, Penny WD, Harrison LM, Daunizeau J, Ruff CC, Duzel E, et al. Integrated Bayesian models of learning and decision making for saccadic eye movements. Neural Networks. 2008;21: 1247–1260. doi:10.1016/j.neunet.2008.08.007

23. Munoz DP. Commentary: Saccadic eye movements: Overview of neural circuitry. Progress in Brain Research. Elsevier; 2002. pp. 89–96. doi:10.1016/S0079-6123(02)40044-1

24. Wurtz RH. Brain mechanisms for active vision. Daedalus. 2015;144: 10–21. doi:10.1162/DAED_a_00314

25. Carpenter RHS, Williams MLL. Neural computation of log likelihood in control of saccadic eye movements. Nature. 1995;377: 59–62. doi:10.1038/377059a0

26. Parr T, Friston KJ. The active construction of the visual world. Neuropsychologia. Elsevier Ltd; 2017. pp. 92–101. doi:10.1016/j.neuropsychologia.2017.08.003

27. Brainard DH. The Psychophysics Toolbox. Spat Vis. 1997;10: 433–436. doi:10.1163/156856897X00357

28. Cornelissen FW, Peters EM, Palmer J. The Eyelink Toolbox: Eye tracking with MATLAB and the Psychophysics Toolbox. Behav Res Methods, Instruments, Comput. 2002;34: 613–617. doi:10.3758/BF03195489

29. Manohar SG. Matlib: MATLAB tools for plotting, data analysis, eye tracking and experiment design (Public). 2019. doi:10.17605/OSF.IO/VMABG

30. Burnham KP, Anderson DR. Multimodel Inference. Sociol Methods Res. 2004;33: 261–304. doi:10.1177/0049124104268644

